# Transcription disrupts DNA-scaffolded bacteriophage repressor complexes

**DOI:** 10.1101/2021.10.29.466366

**Authors:** Yue Lu, Zsuzsanna Voros, Gustavo Borjas, Cristin Hendrickson, Keith Shearwin, David Dunlap, Laura Finzi

## Abstract

DNA can act as a scaffold for the cooperative binding of protein oligomers. For example, the phage 186 CI repressor forms a wheel of seven dimers wrapped in DNA carrying specific binding sites, while the phage λ CI repressor binds in units of dimers to two well-separated sets of operators, forming a DNA loop. Atomic force microscopy was used to measure transcription elongation by *E. coli* RNA polymerase through these protein complexes. 186 CI or λ CI bound along unlooped DNA negligibly interfered with transcription by RNAP. More complex topologies induced by scaffolded, cooperatively bound repressor oligomers did not form significantly better roadblocks to transcription. Thus, despite binding with rather high affinity, these repressors are not effective roadblocks to transcription.

## Introduction

Genomic DNA is decorated with proteins which may have various and/or multiple functions. Proteins may compact DNA [1, 2], protect it from the damaging effect of external agents [3, 4], or mediate long-range interactions implicated in processes such as the management of DNA superhelicity [5, 6], repair [7–9], or transcription regulation [10, 11]. Whatever their function(s), these proteins often are found on the path of transcription elongation complexes and may impede the processivity of RNA polymerase (RNAP). The mechanisms by which RNAP might overcome such roadblocks are poorly understood. Single-molecule approaches are ideally suited to dissect such mechanisms and previous studies have focused on the ability of RNA polymerase to disrupt nucleosomes [12–15]. However, the histone proteins which form nucleosomes interact with DNA non-specifically and are substrates for extensive post-translational and epigenetic modifications that regulate chromatin remodeling and the recruitment of accessory factors that regulate transcription [16, 17]. In contrast, simpler transcription factors (TFs) from organisms spanning all kingdoms recognize specific sites on DNA [18, 19] to shape the genome and regulate various genomic functions without chemical modifications regulated by complex pathways. These TFs may recognize several specific sequences to which they bind cooperatively with different affinities [20, 21]. Such cooperative interactions may produce topology such as wrapped or looped DNA, which have not been directly investigated in earlier studies on transcription through roadblocks either *in vivo* or *in vitro.*

Recently, it was shown *in vitro* that an advancing *E. coli* RNAP on a linear template is effectively blocked by the lac repressor (LacI) protein bound to strong operators, such as Oid or O1 (Kd = 0.13 nM and 0.6 nM, respectively) but not to the weaker O2 (Kd = 2.7 nM) [22]. Lac repressor is a homo-tetramer with two dimeric DNA binding units and can bind two operators simultaneously, securing a loop in the intervening DNA [23, 24]. Remarkably, even LacI bound to O2 significantly impedes transcription when it is part of a loop [25, 26]. Many transcription factors secure loops between distant sites [27, 28], and others may wrap DNA [29]. Here, we extended our study of topological impediments of transcription elongation to include the 186 and the λ CI repressors, two cooperatively binding proteins with packaging and regulatory functions that, respectively, wrap or loop DNA [29, 30]. After bacteriophage 186 infection of *E. coli,* the 186 CI repressor maintains lysogeny via repression of lytic genes [31, 32]. Dimers of 186 CI oligomerize via their C-terminal domains to form a wheel-shaped heptamer around which DNA can wrap as three of the seven dimers bind to three strong (K_D_ = 30 nM) [31], specific binding sites at the lytic pR promoter, favoring weaker binding of other dimer spokes of the wheel to adjacent non-specific sequences which include pL [29, 33, 34] (Figure S1A). Two other specific binding sites, FL and FR, are found, each on either side of pR. When pL is not wrapped on the wheel, the latter may bridge pR and either FR or FL [35] (Figure S1B). In this study we used a template containing the pR/pL and FR sequences to measure interference by 186 CI protein assembly-mediated DNA wraps and loops with transcription elongation.

The λ CI repressor has a similar role to that of 186 CI in maintaining lysogeny after λ bacteriophage infects *E. coli* [30, 36, 37]. The λ repressor binds as a dimer to each of three adjacent binding sites (OL1, OL2 and OL3) separated by 2.3 kbp from another similar set (OR1, OR2 and OR3) [38] (Figure S1C). The N-terminal domains of dimers bind DNA, leaving the C-terminal domains to interact with an adjacent or juxtaposed dimer [39, 40]. These cooperative protein-protein interactions stabilize a DNA loop between the OL and OR regions [41–46] (Figure S1D). To test interference with transcription elongation by these different protein mediated-DNA topologies, the progress of RNAP was measured on DNA templates in which the 186 CI could mediate either a DNA wrap or loop, as well as on templates where λ CI could mediate a loop. We analyzed snapshots of transcription elongation complexes imaged with the atomic force microscope (AFM) similarly to that reported in a recent study of the lac repressor protein [47]. Neither the 186 nor the λ CI repressors were strong roadblocks, regardless of the DNA topology induced. While the LacI protein on a high affinity binding site or in a looped confirmation blocks transcription [47], the cooperative proteins assemblies studied here do not constitute significant roadblocks to transcription. Since proteins, such as CTCF in eukaryotes, or HU in prokaryotes also circumscribe large loops of transcriptionally active domains (TADs) [48–50], it is intriguing to think that assemblies of architectural protein may become roadblocks only when forming much higher-level aggregates.

## Material and Methods

### DNA constructs for scanning force microscopy

The DNA templates used for AFM measurements of transcription through 186 CI obstacles were 1510 bp in length and were produced by PCR using plasmid template pT7A1_pRpL-FR (see supplementary information), with an unlabeled forward primer (5’-AGCCATGACCCAGTCAC) and a 5’-biotin-labeled reverse primer (5’-GCACTCTCAGTACAATCTGCTCTG). PCR products were purified using a GeneJet PCR Cleanup kit (Thermo Scientific, Waltham, MA). From the forward primer the fragments contained the T7A1 promoter with a transcription start site 167 bp, followed by the three 186 pR/pL binding sites 271 bp further, the FR site at 391 bp beyond that, and finally a λt1 terminator at 551 bp beyond FR (Figure 1A).

**Figure 1.**
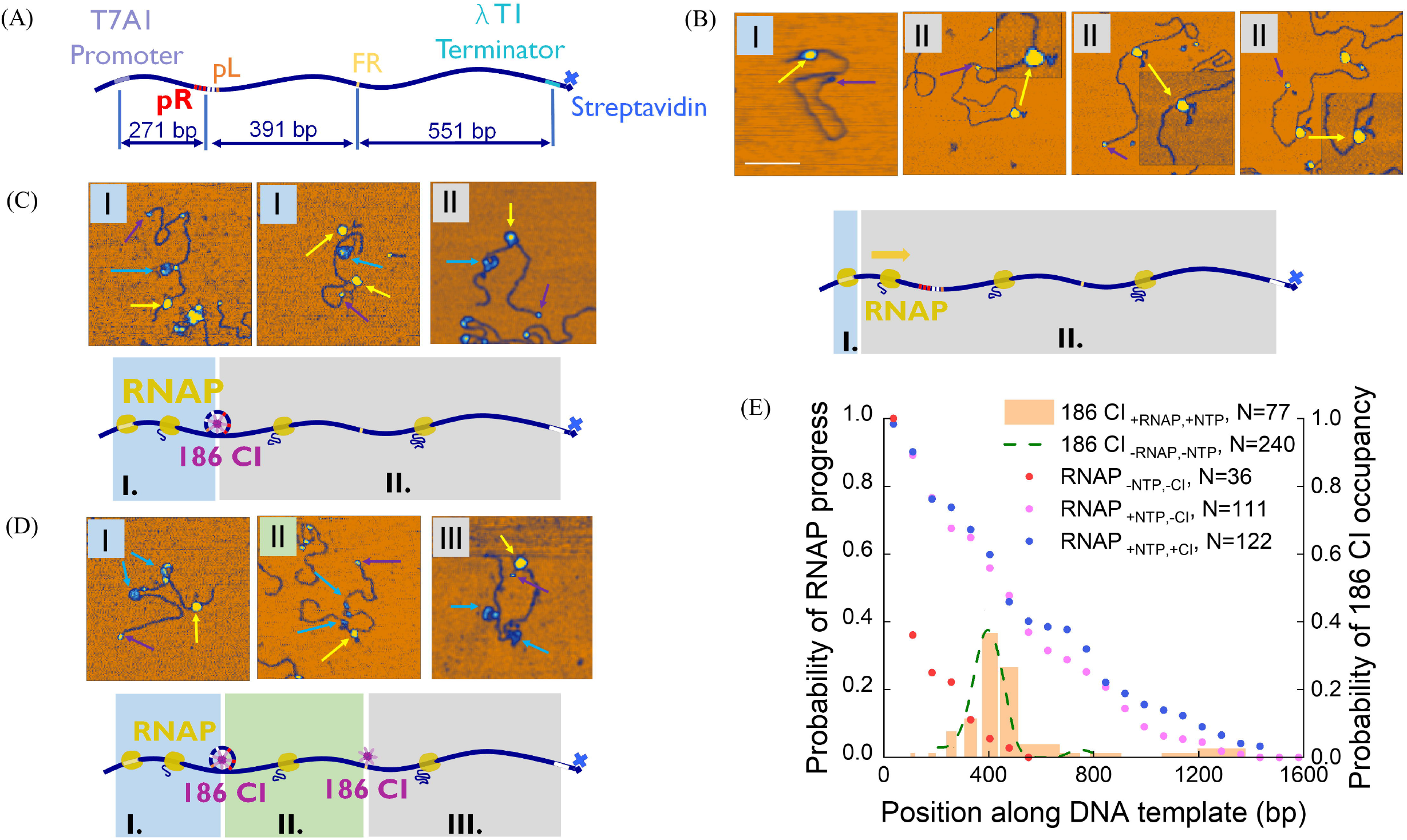
RNAP elongation in the presence of the 186 CI repressor observed using AFM. **(A)** A diagram of the DNA template shows the spacings between relevant DNA features. **(B)** The upper panel shows AFM images of control experiments in the absence of 186 CI, and the lower panel is a color-coded cartoon illustrating categories of RNAP progress. RNAP particles are marked by yellow arrows and streptavidin are smaller particles marked by purple arrows. Nascent RNA associated with all RNAP particles is visible as dark blue blebs on the periphery of the particles. **(C)** The upper panel is a gallery of transcription elongation complexes on DNA templates wrapped around 186 CI bound at pR, and the lower panel is a color-coded cartoon illustrating categories of RNAP progress. 186 CI particles are marked by cyan arrows. **(D)** The upper panel is a gallery of transcription elongation complexes on DNA templates wrapped around 186 CI bound at pR as well as a repressor bound at FR, and the lower panel is a color-coded cartoon illustrating categories of RNAP progress. **(E)** The probability of progress by RNAP is plotted for different experimental conditions with the measured positions of 186 CI indicated as a bar histogram. The colored dots indicate the fraction of elongation complexes found at distances greater than or equal to *x* from the promoter along the DNA molecule. Heights in AFM images rise from orange color-coded low values to dark blue to cyan and the highest features appear yellow.

The DNA templates used for AFM measurements of transcription through λ CI obstacles were 1737 bp in length and were produced by PCR using plasmid template pDK_LMB_400 (see supplementary information), with unlabeled forward (5’-CTTGTCTGTAAGCGGATGCC) and reverse (5’-ACGCAAACCGCCTCTCC) primers and purified using a GeneJet PCR Cleanup kit (Thermo Scientific, Waltham, MA). From the forward primer the templates contained the T7A1 promoter with a transcription start site 128 bp, followed by the OL region 261 bp further, the OR region 451 bp beyond that, and finally a λt1 terminator at 678 bp beyond OR (see supplementary information).

### Sample preparation for scanning force microscopy

A 10 μL droplet of 0.01 μg/ml of poly-L-ornithine (1 kDa Mw, Sigma-Aldrich, St. Louis, MO) was deposited onto freshly cleaved mica and incubated for 2 min. The poly-L-ornithine-coated mica was then rinsed drop-wise with 600 μl of high-performance liquid chromatography-grade water and dried with compressed air. Nucleoprotein complexes of RNA polymerase/186 CI, or λ CI, bound at the promoter (promoter complex: PC)/CI binding sites were produced by incubating 1 nM of DNA with 250/300/500 nM 186 CI and 0.1 μM streptavidin, or 150 nM λ CI, and RNA polymerase holoenzyme (New England Biolabs, Ipswich, MA) diluted in transcription buffer (TXB: 20 mM Tris-glutamate (pH 8.0), 10 mM magnesium-glutamate, 50 mM potassium-glutamate, 1 mM DTT) to a final concentration of 6 U/ml for 30 min at 37 °C. To initiate transcription, the reaction mixture was spiked with 1 mM NTPs to give a final NTP concentration at 100 μM, and incubated at 37 °C for 2 min. Then, 250 mM EDTA in TXB was added to give a final concentration of 20 mM EDTA and incubated at 37 °C for 30 s to terminate transcription. 5 μl of this final mixture was deposited on the poly-L-ornithine-coated mica and incubated for 2 min. This droplet was rinsed with 600 μl of high-performance liquid chromatography-grade water and dried gently with compressed air [35, 47, 51].

### Scanning force microscopy and DNA/protein data measurement

Images were acquired with a NanoScope MultiMode VIII AFM microscope (Bruker Nano Surfaces, Santa Barbara, CA, USA) operating in soft tapping mode, or ScanAsyst^®^ mode, using cantilevers with 2 nm nominal tip radius. Areas of 4×4 μm^2^ were scanned at a rate of 0.5 Hz with a resolution of 2048×2048 pixels. After filtering the images to remove scan line offsets, tilt, and bow, DNA/protein molecules were measured with NanoScope Analysis or the NeuronJ plugin [52] of ImageJ [53]. The length measurement function was used to obtain the contour length of DNA length and establish the position of particle binding; lengths were then normalized by setting them all to 1510 bp for the 186 CI binding fragment, or 1737 bp for the lambda CI binding fragment. The Particle Analysis function of the NanoScope Analysis software was used to measure the diameter, height and volume of bound protein molecules.

## Results

### 186 CI negligibly blocks transcription

#### 186 CI and RNA polymerase can be distinguished by height

Scanning force microscopy was used to determine the disposition of RNAP transcribing through 186 CI – DNA nucleoprotein obstacles. Micrographs of transcription elongation complexes in the presence of the 186 CI repressor were acquired to score the location of RNA polymerase and the 186 CI obstacle along DNA templates that contained two 186 CI-binding regions. The first binding region contained the three 186 strong pR sites 271 bp downstream of the T7A1 promoter and adjacent to the weaker pL site. Together these sites can wrap around the 186 heptamer to form a stable wheel-like structure. The second site, FR, which has a lower affinity for 186 CI than pR, was located 391 bp downstream of pR. A λt1 terminator was located 551 bp downstream of FR and the 5’ strand of this end of the template was labeled with a biotin [Figure 1A]. 186 CI (500 nM), streptavidin and RNAP were added to a DNA solution at the same time and transcription was activated and stalled as described in Materials and Methods by addition of NTPs and EDTA, respectively. An aliquot of the sample was then immediately deposited on poly-L-ornithine-coated mica.

Streptavidin bound to the biotin label marked the terminator end of the DNA template (Figures 1B, C, & D and S2A, purple arrows). The majority of the DNA molecules had RNAP bound either at the promoter or associated with nascent RNA along the DNA template (Figures 1B, C, & D and S2A, yellow arrows), and 186 CI bound at the pR/pL and/or the FR site (Figures 1C & D and S2A, cyan arrows). Circular 186 CI and RNAP particles in AFM images had similar diameters ranging 15 – 25 nm. While RNA polymerase averaged 4.5 nm in height, 186 CI averaged only 2 nm (Figure S2C). Thus, 186 CI and RNAP were easily distinguished in AFM micrographs.

#### 186 CI does not block transcription

To illustrate the analysis and interpretation of the AFM images, nucleoprotein complexes were categorized based on the position of RNAP along the DNA template in different experimental conditions. In measurements in the absence of 186 CI, we found some RNAP molecules at promoters and others that had transcribed some distance (Figure 1B upper panel) and categorized them as shown in the lower panel of Figure 1B. Note that nascent RNA is clearly visible associated with elongating RNAP in all images as highlighted in the enlarged areas shown in Figure 1B.

Adding 186 CI before activating transcription produced a majority of repressor complexes bound to the pR/pL site only (Figure 1C) and a small number bound to both the pR/pL and FR sites (Figure 1D). However, loops between pR/pL and FR were not observed and indeed they were estimated to be rare in a previous study [54]. In a few cases (12%), 186 CI was found only at FR (Figure S3). We classified the nucleoprotein complexes according to the position of RNAP with respect to the pR/pL and FR binding sites (Figure 1C & D lower panels). RNAP was found in all segments of the DNA template. To better summarize the data, the positions of RNAP along the DNA were binned and the fraction of elongation complexes found at or beyond each bin was plotted in each experimental condition (Figure 1E). These plots indicate that the presence of the 186 CI wheel does not affect RNAP progress.

#### Transcribing RNA polymerase may cause the dissociation of 186 CI

In images like those in Figure 1, 186 CI particles behind RNAP (Figures 1C image II, 1D images II & III) often appear smaller than those ahead of RNAP (Figures 1C image I, 1D images I & II). To test the idea that RNAP transcribing through the heptamer may cause the dissociation of some of its dimers from the wheel, we measured the diameter (Figure 2A) and volume (Figure 2B) of 186 CI under different conditions of transcription. In control experiments with 186 CI and with or without added RNAP but no NTP, the mean diameter was 16-17 nm with a wide distribution as previously reported [35]. The mean volume was about 650 cubic nanometers. Non-translocating RNAP did not affect the size of 186 CI. Diameters of RNA polymerase were more narrowly distributed than those of 186 CI (Figure S4), which indicates labile 186 CI oligomerization. Following a minute of transcription, there was a slight decrease in both the diameter and volume of the 186 CI repressor. A Welch’s unequal variances t-test of the data with and without transcription (NTP) produced *t* equal to 5.24 for 102 degrees of freedom, corresponding to a *p* value smaller than 0.0001. This confirms transcription significantly reduced the volume of 186 CI particles.

**Figure 2.**
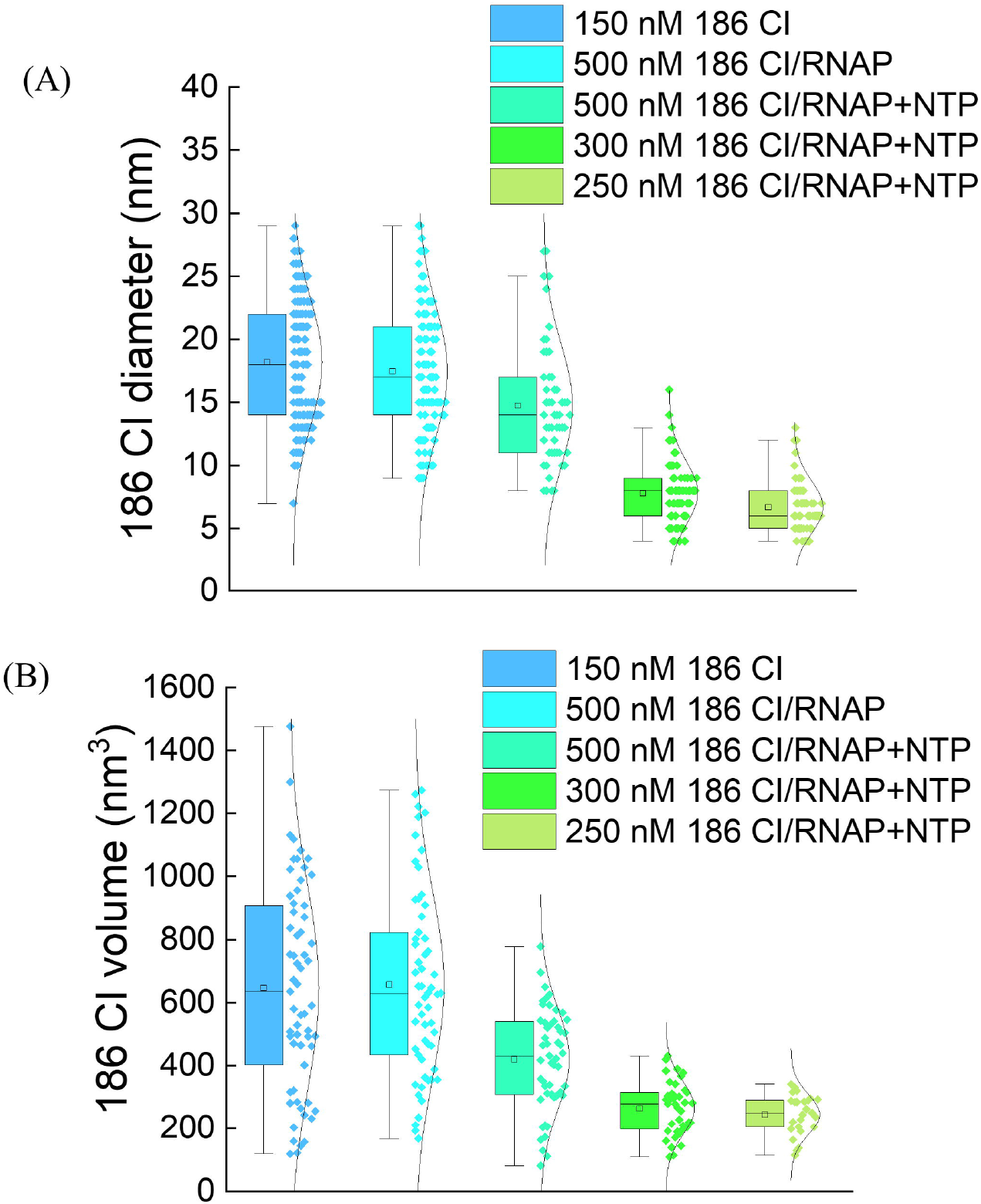
The 186 CI heptameric wheel is disrupted by transcription. Measurements of the diameters **(A)** and volumes **(B)** of 186 CI particles measured in AFM transcription assays are shown in whisker and violin plots. The conditions from left to right are: +150 nM 186 CI/-RNAP/-NTP; +500 nM 186 CI/+RNAP/-NTP; +500 nM 186 CI/RNAP/+NTP; +300 nM 186 CI/+RNAP/+NTP; +250 nM 186 CI /RNAP/+NTP.

Figure 2 shows that the volume of 186 CI is largest, about 650 nm^3^, in the absence of transcription. This volume is consistent with what previously found for the full 186 CI wheel composed of seven dimers [35]. Decreasing the concentration of 186 CI from 500 to 300 nM in conditions of transcription, further decreased of the volume of 186 CI particles to ~420 nm^3^ which would correspond to 4 – 5 dimers. The mean volume of 255 nm^3^ measured at the lowest 186 CI concentration used, 250 nM. This value is consistent with a 186 CI complex containing 3 dimers. Thus (i) advancing RNAP may partially disrupt the 186 CI wheel, and (ii) after disruption 186 CI appears to re-associate with the DNA in a concentration dependent manner.

### λ CI mediating a DNA loop weakly obstructs transcription

#### The composition, conformation and elongation stage of transcription complexes were easily identifiable in AFM micrographs

The DNA construct used in AFM measurements of transcription through the λ CI repressor contained two sets of three operators for the λ CI protein. The OL region, located 261 bp downstream of the T7A1 promoter contained sequentially the OL1, OL2 and OL3 operators. The OR region containing sequentially the OR3, OR2, and OR1 operators was 451 bp beyond OL and was followed by the lambda T1 terminator 678 bp further on (Figure 3A). Due to the relative binding affinity of the operators for λ CI, OR1≈OR2≈OL1≈OL2>OL3>OR3 [55, 56], and the cooperative interaction of adjacent λ CI dimers [39], one might expect incomplete occupancy of all binding sites on linear DNA [46, 57]. However, looping stabilizes occupancy of λ CI dimers at all sites both *in vitro* [42] and *in vivo* [46, 57]. As in the experiments described for 186 CI, transcription was activated and halted by adding NTP and EDTA, respectively (Materials and Methods) and a small aliquot of the sample was immediately deposited on poly-L-ornithine-coated mica for imaging.

**Figure 3.**
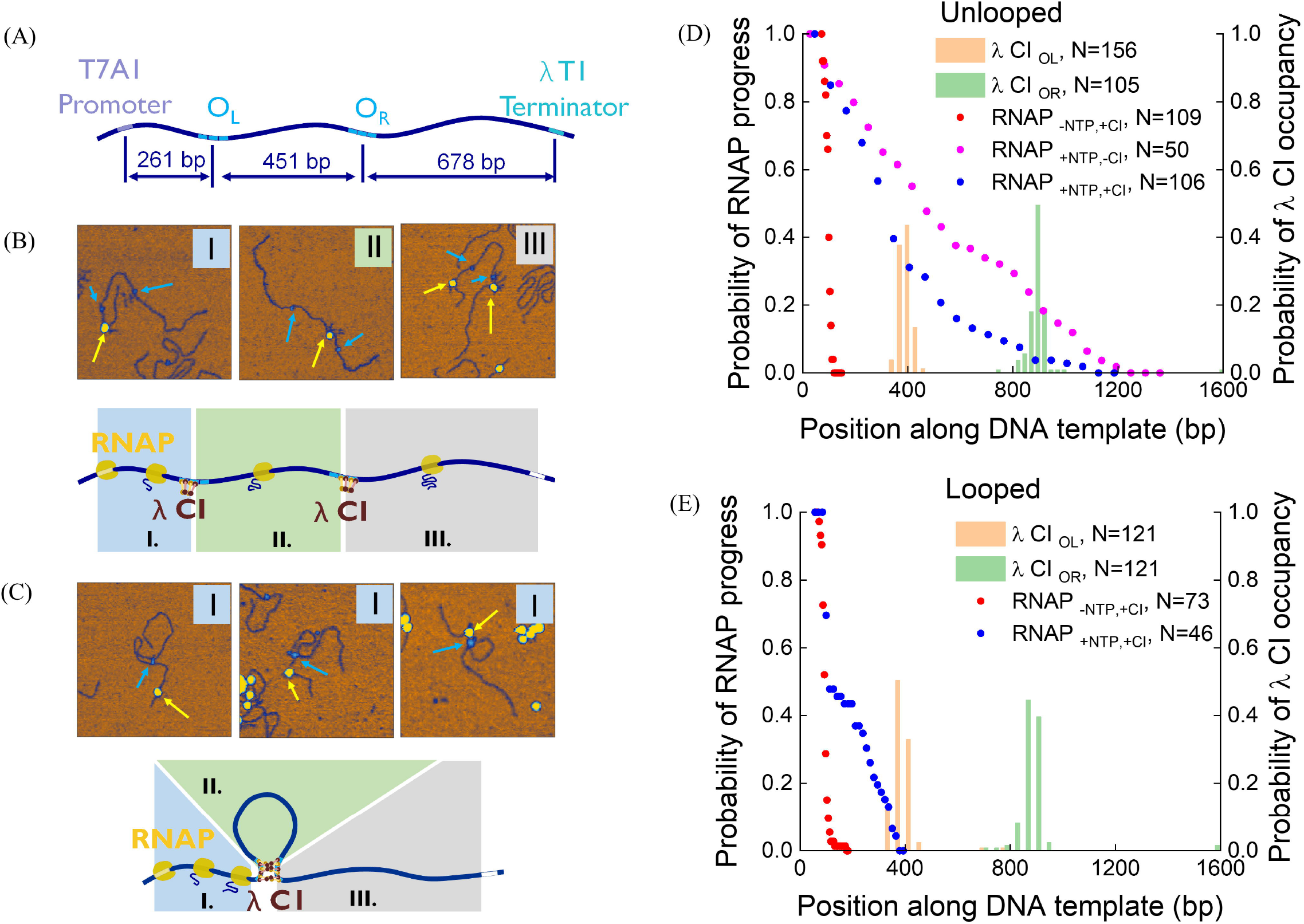
RNAP elongation in the presence of lambda CI repressor. (Left) **(A)** A diagram of the DNA template shows the spacings between relevant DNA features. **(B)** The upper panel shows transcription elongation complexes along a linear DNA template containing the OL and OR lambda operators in the presence of the lambda CI, and below is a color-coded cartoon illustrating categories of RNAP progress. **(C)** The upper panel shows representative transcription elongation complexes along a looped DNA template, and below is a color-coded cartoon illustrating categories of RNAP progress. Colored arrows marking features are as in Figure 1. The top right AFM image includes a second RNAP that had started elongation of the same DNA template, following a first RNAP that had just bypassed λ CI bound at the OR region before stalling. No looped molecules were found with RNAP inside or after the loop. Heights in AFM images rise from orange color-coded low values to dark blue to cyan and the highest features appear yellow. (Right) Plots of the probability of RNAP progress in the presence of the lambda CI repressor in either unlooped **(D)** or looped **(E)** DNA templates. The bar histogram indicates the location of lambda CI bound at the OL (orange) and OR (green) binding sites. The colored dots indicate the fraction of elongation complexes found at distances greater than or equal to *x* from the promoter along the DNA molecule. Heights in AFM images rise from orange color-coded low values to dark blue to cyan and the highest features appear yellow.

AFM images of transcription complexes were recorded and the positions of RNAP and λ CI protein along the unlooped/looped templates were tabulated. RNAP (MW ~450 kDa) is much larger and easily distinguished from even a λ CI dodecamer (MW_dimer_ ~ 52 kDa; Figure 3B & C). Although some unlooped DNA molecules had λ CI bound to only the OL or the OR region, most of the DNA molecules had both OR and OL regions occupied by λ CI. OL and OR were distinguished by their positions with respect to the closest end: ~300 bp for OL and ~600 bp for OR. Figure 3B&C shows representative images of transcriptional complexes on unlooped and looped DDNA templates, accompanied by cartoon describing the progress of RNAP. The unlooped complexes were classified according to the position of RNAP before OL, between OL and OR, or after OR. The looped complexes were scored according to their position before, inside, or after the λ CI-mediated loop.

#### Unlooped DNA-λ CI complexes do not impede transcription elongation

Upon addition of NTPs, transcribing RNAP was found throughout the length of unlooped DNA templates indicating that λ CI is an insignificant obstacle for RNAP. In addition, most molecules exhibited λ CI bound to operators both ahead and behind RNAP (Figure 3B) suggesting that λ CI promptly rebinds to operators after having been dislodged by RNAP.

The cumulative progress of transcription elongation complexes and the positions of λ CI oligomers along the DNA molecules is shown in Figure 3D. The frequency distribution of the positions λ CI along the DNA template is shown as the orange and green bar histogram, which demonstrates specific binding. The red data points represent cumulative RNAP progress without NTP. In this condition, RNAP is only found at the promoter (Figure 3D; Figure S5, left). The pink curve represents the progress of RNAP with NTP but without λ CI (Figure 3D; Figure S5, right). The blue curve represents transcription progress in the presence of λ CI and shows a somewhat steeper decline, perhaps reflecting that λ CI weakly interferes with RNAP. If λ CI were a strong obstacle for the transcription elongation complex, the blue curve in Figure 3D would be expected to display a steep drop in the probability of finding RNAP at, or past, the OL region.

#### λ CI mediating a DNA loop does not substantially impede transcription elongation

In all templates in which λ CI mediated a DNA loop, transcription elongation complexes (TECs) were found before but not within or after the loop (Figure 3C, 3E). As for the 186 CI repressor, the frequency distribution of λ CI positions along the DNA templates shown as the orange and green bar histogram which indicates that λ CI binds OL/OR specifically. The red curve represents RNAP progress without NTP and all TECs clustered at the promoter (Figure S5, left). The blue curve, instead, represents RNAP progress on DNA looped by a λ CI oligomer. The fraction of observed TECs decreases rapidly even before the first λ CI peak at the OL region. This observation might seem to indicate that RNAP polymerase did not pass the looped DNA-λ CI complex and that impediment by the λ CI roadblock in a loop increases considerably. However, when the DNA-binding probability in different experimental conditions was plotted for the λ CI repressor (Figure 4B), it became apparent that transcription (blue bars) dramatically reduced the number of λ CI-mediated loops, and indeed of DNA-λ CI complexes all together, casting doubt on the roadblocking ability of the λ CI protein. A similar, if less pronounced, effect was seen in the case of DNA-186 CI complexes (Figure 4A),

**Figure 4.**
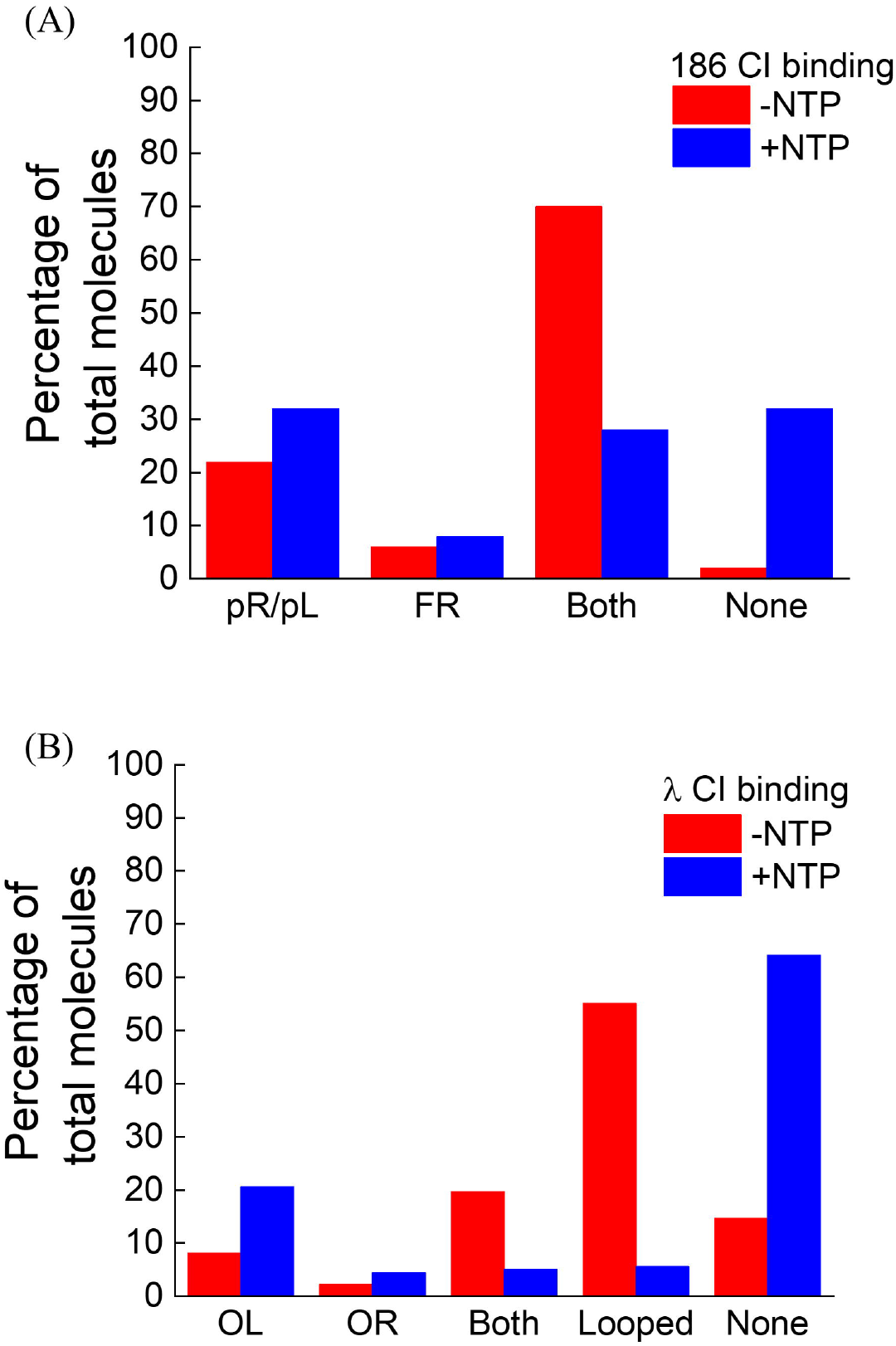
Transcription reduces the number of repressor molecules bound to DNA. These plots show the numbers of different configurations of DNA molecules bound by λ CI **(A)** or 186 CI **(B)** particles with (red) or without (blue) transcription. (A) Actively transcribing RNAP decreased the number of λ CI-mediated looped complexes substantially, while the number of DNA molecules with no repressor molecules bound increased compared to the number observed in the absence of NTPs. (B) An increased proportion of 186 DNA molecules with no 186 CI bound, and a decreased proportion of DNA molecules with both pR and FR bound by CI were observed in conditions of active transcription.

In the case of the λ CI repressor (Figure 4B), in the absence of transcription about 15 % of all DNA molecules analyzed did not have any repressor bound to them. However, transcription increased that percentage to 64 %. In addition, without transcription about 10 % of the DNA molecules with λ CI had only one operator region (either OL or OR) occupied, which is low compared to the 75 % with λ CI at both OL and OR (including unlooped and looped DNA-λ CI complexes without transcription). With transcription, the percentage of DNA molecules with λ CI bound either at OL or at OR increased to 25 %, while the percentage of DNA molecules with both regions occupied by λ CI (unlooped and looped) dropped to 10 %. Before transcription, the percentage of DNA molecules looped by λ CI was 55 %, while the unlooped nucleoprotein complexes were 20 %, but both decreased to just 5 % after transcription. These decreases suggest that transcribing RNAP induces the dissociation of λ CI from DNA. The slight increases in the number of DNA molecules bound by only one λ CI oligomer indicate that some of the dislodged λ CI may reassociate after RNAP passes through the operator sites.

In the case of the 186 CI repressor (Figure 4A), in the absence of transcription, nearly all DNA molecules had bound repressor with only 2 % of DNA molecules without any 186CI. In the presence of transcription, the percentage of DNA molecules without repressor increased to 32 %. Similarly, the percentage of DNA molecules with both pR/pL and FR bound by the 186 CI repressor is 70 % in the absence of transcription but only 28 % in the presence of transcription. Curiously, before transcription, the percentages of 186 CI repressor bound to either pR/pL or FR were 22 % and 6 %, respectively. This difference is consistent with the differences in affinity of the repressor for these two binding sites [31]. In the presence of transcription, these percentages increased slightly to 32 % and 8 %, respectively.

## Discussion

The λ CI and 186 CI repressors cooperatively assemble on DNA binding sites where they may block transcription by RNAP. The temperate λ and P2-like 186 bacteriophages are very different, belonging to two different phage families, yet their CI repressors have some structural similarities [58, 59]. Their N-terminal domains bind to DNA via a helix-turn-helix motif, while the C-terminal domains mediate both dimer formation and higher order structures. Proteinprotein cooperativity thus plays an important role in the regulatory function of these two repressors, permitting formation of topological forms such as DNA loops or wraps.

In contrast to the lac repressor protein, which binds DNA as a bidentate tetramer and forms a strong roadblock to transcription, neither the λ CI nor the 186 CI repressor proved to be significant roadblocks to RNAP. Although a plot of the surviving fraction of elongation complexes as a function of distance from the promoter along looped DNA molecules (Figure 3E) drops to zero at OL as if the λ CI repressor were a strong roadblock, comparison of λ CI DNA-binding probability in the absence and presence of transcription, and comparison with the corresponding DNA-binding probability graphs for the 186 CI protein (Figure 4B) and the lac repressor (Figure S4 in [47]) do not support this interpretation.

The analysis of AFM micrographs shows that although λ CI mediating a DNA loop appeared to block transcription efficiently (Figure 3E), the number of looped molecules was much lower than in the absence of transcription (Figure 4B), indicating that transcription either interferes with loop formation and/or that RNAP disrupts the loop and transcribes past. Thus, λ CI is not a strong roadblock even when mediating a DNA loop. This is in agreement with results from *in vivo* experiments [60].

The 186 CI repressor dimers oligomerize to form a wheel of seven units, which preferentially wraps DNA at pR/pL. This wheel was not a roadblock to an elongating RNAP. The 186 CI repressor can also bridge pR and FR (or FL) binding sites inducing a DNA loop [34, 54], although it occurs with a low probability *in vitro* [35] and was never observed in the transcription measurements reported here. The lack of such loops in AFM images of DNA templates with active transcription elongation complexes indicates that they are easily disrupted by elongating RNAPs.

Analysis of the transcription templates also indicates that the λ and 186 CI repressors may reassociate with the DNA template after being dislodged by RNAP. 186 CI may rebind more easily than λ CI, albeit in smaller oligomers, but a lambda loop rarely reformed after RNAP passage (Figure 4B).

A possible mechanism by which RNAP may bypass each repressor is illustrated in Figure 5. An elongating RNAP likely causes the dissociation of successive dimers in the wheel and when the three dimers bound to the highest affinity pR sites are dissociated, the whole wheel may disperse. Based on the volumes of particles measured in Figure 2, the most likely particles to rebind will consist of three dimers, consistent with cooperative binding to the three strong operators at 186 pR/pL. This nucleation might eventually recruit more dimers, until the full 7-dimer wheel has reassembled. The rate of repressor reassociation after passage of an RNAP, relative to the rate of transcription, has been shown to be an important factor in studies of transcriptional interference [60].

**Figure 5.**
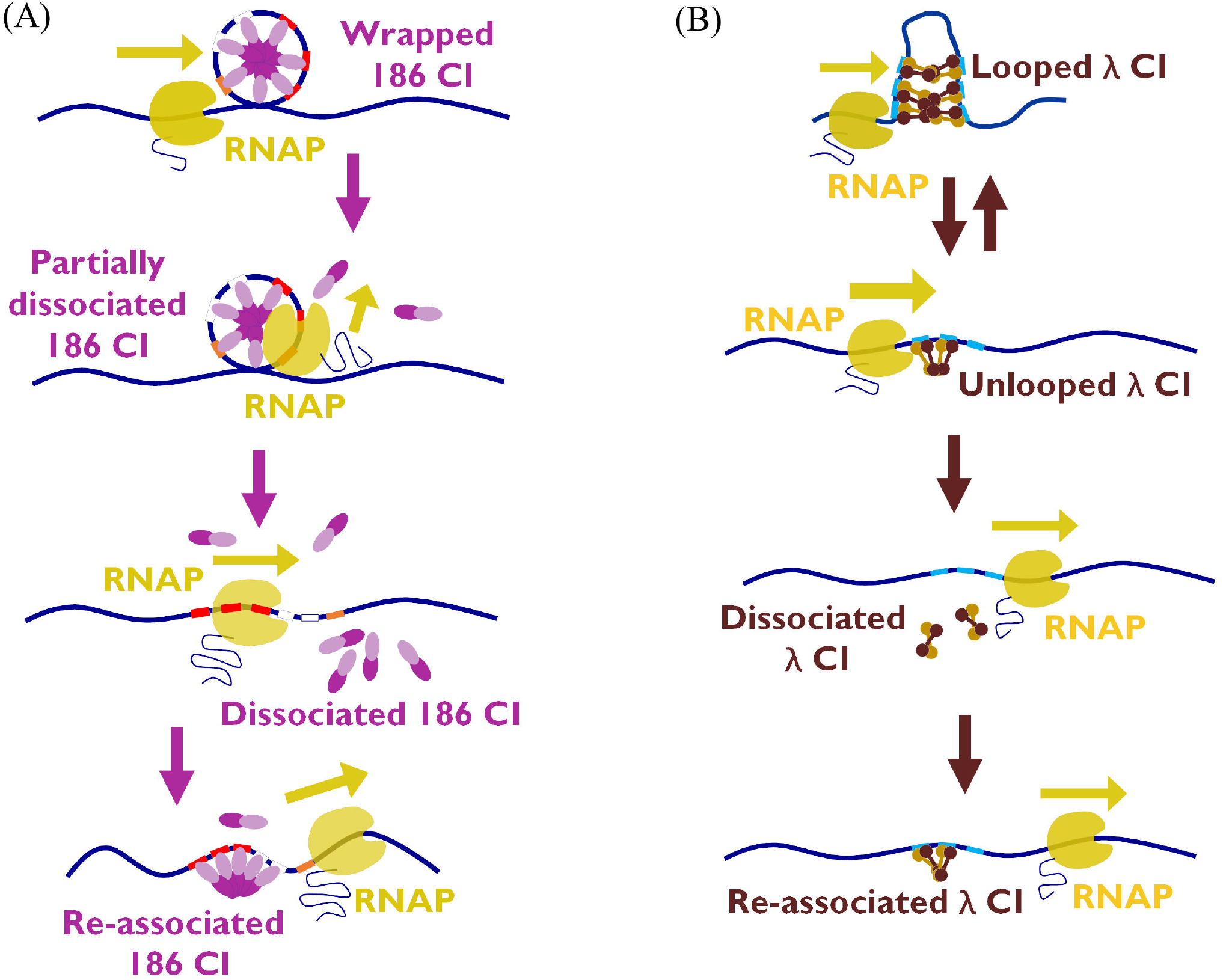
Schematic of RNA polymerase disrupting the 186 CI **(A)** and λ CI **(B)** repressors. RNA polymerase likely disrupts the 186 CI complex causing the wheel-like structure to dissociate, but 186 CI may reassemble on the high affinity PR sites after RNAP passage. For λ CI, disruption of the first bound dimer encountered by advancing RNAP, and loss of the cooperative interactions between dimers, may weaken binding by subsequent dimers, allowing them to be more easily dislodged. After RNAP passage, the dimers may slowly reassemble on the DNA.

Figure 5B depicts the case of the λ CI repressor. λ CI closing a DNA loop is a weak roadblock for RNAP; when the λ Ci-mediated loop breaks down, either by spontaneous dissociation or as the result of active dislodging by RNAP, RNAP causes the dissociation of the λ CI dimers from their operators and may also disrupt any oligomers that might have formed. After passage of the RNAP, λ CI dimers can rebind DNA.

In conclusion, the present study shows that the λ CI and 186 CI repressors do not interfere significantly with transcription elongation complexes and suggests that cooperative binding is a critical feature which allows these labile repressors to quickly regain their regulatory function on the DNA template. Figure 6 graphically summarizes the results for the proteins and operator patterns tested. Proteins binding to stable, high affinity binding sites block transcription most effectively. Although cooperative protein assemblies may induce reinforcing topologies like wrapping or looping that potentiate intermediate affinity sites like the lac repressor O2 site [25], they are generally weaker roadblocks. It may be that weak roadblocking of RNAP *in vivo* is crucial to maintain viability, as shown by experiments using a very tight binding mutant of the EcoRI protein which, if inserted at different loci in the genome is lethal to *E. coli* [61].

**Figure 6.**
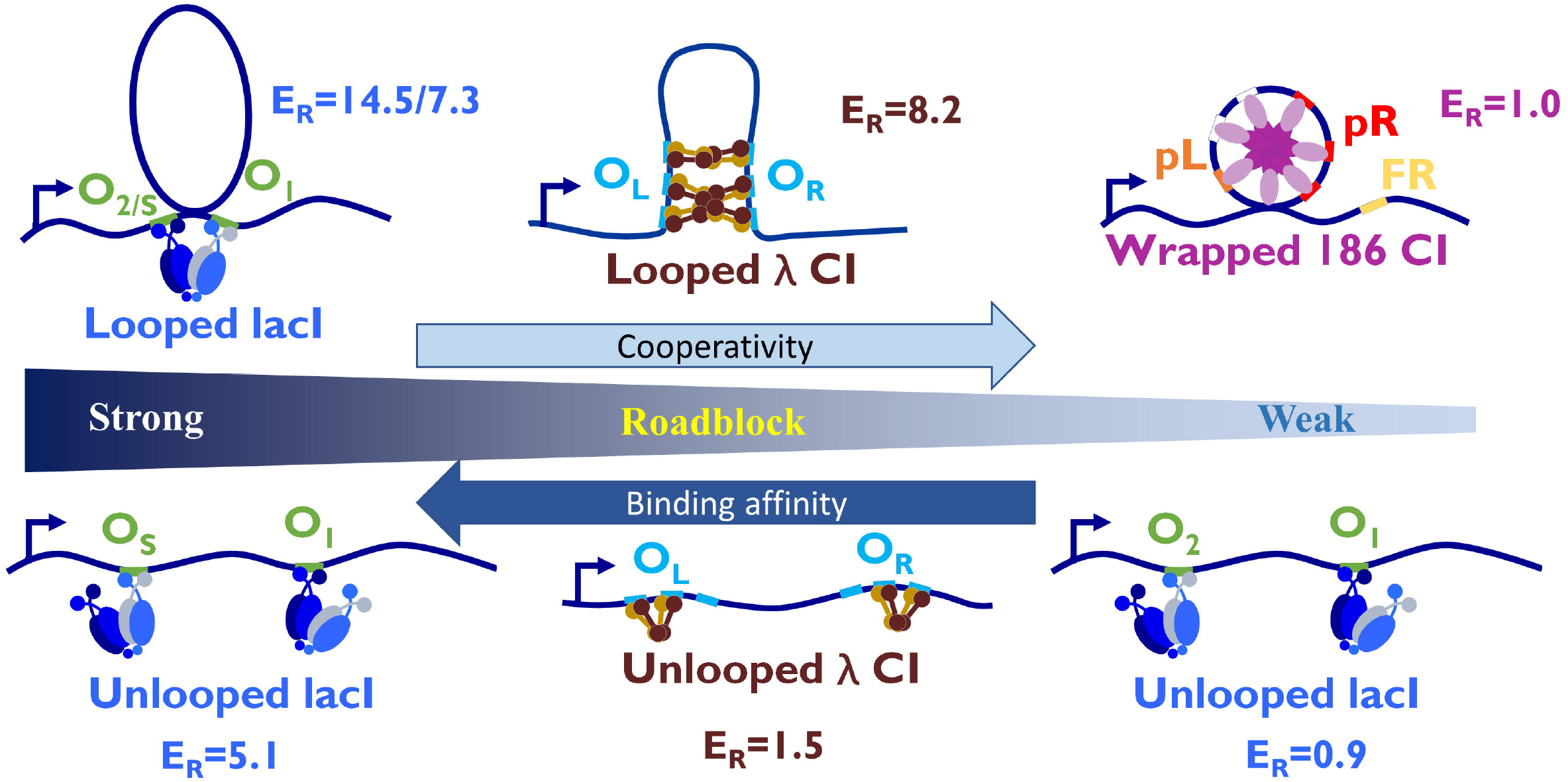
A ranking of the potential roadblock proteins tested emphasizes that proteins binding to stable, high affinity binding sites block transcription most effectively (SI). Lac repressor binding to Os or looped O2 was most effective. Cooperative λ CI oligomers were intermediate strength roadblocks (Table S1), while cooperative binding to lower affinity sites as in the 186 PR sequence, did not impede RNAP significantly.

## Acknowledgments

This work was supported by the National Institutes of Health to L.F. (R01 GM084070) and Australian Research Council to KS (DP150103009). Kathleen Matthews at Rice University generously provided the LacI protein. We thank Allison Cartee for help measuring RNAP locations on templates in the presence of 186 CI and Ian Dodd for comments on the manuscript.

