## Supplementary material for "Transcription disrupts DNA-scaffolded bacteriophage repressor complexes": Figures S1-5, Table S1 and SI

**Quantifying roadblock strength**

First, the CI repressor position is identified by the peak of the frequency distribution histogram (Figures 1E and 3 D & E). Then, we define the strength of a roadblock, $E_R$ as follows:

$$E_R = {(P_{b+}/P_{a+})}\Big/{(P_{b-}/P_{a-})}$$

where $P_{b+}$ is the probability of finding an RNAP before the obstacle position in the presence of CI repressor (λ or 186) while $P_{a+}$ is the probability of finding an RNAP after the obstacle; $P_{b-}$ and $P_{a-}$ are the corresponding probabilities in the absence of CI. The position subscripts *b* and *a* indicate the full width, half height positions bracketing the peak of repressor distribution. Thus, a low probability of RNAP traversing a high strength roadblock would give a high $E_R$ score, while a protein which had no ability to block RNAP would have an $E_R$ score of 1.

Below is a table of the probabilities of RNA progress straddling obstacle positions and the associated roadblock strengths calculated for measurements on templates bearing the lac, lambda CI, and 186 CI repressors.

Previously published measurements on the roadblock activity of LacI *in vitro*, considered three topologies: (1) LacI-mediated looped Os-O1 or O2-O1 DNA; (2) Os-O1 unlooped DNA and (3) O2-O1 unlooped DNA, where Os (also known as $O_{ideal}$) is an engineered operator with binding affinity > O1>O2.

For LacI-mediated looped DNA, a loop between the near and far binding sites produced strong roadblocks with $E_R$ = 7.3 and 14.5 for looped $O_s$-$O_1$ and $O_2$-$O_1$ respectively. For unlooped $O_s$-$O_1$ DNA, $E_R$ = 5.7 at the first operator and 0.9 at the second operator. The composite roadblock efficiency of templates including both obstacles, can be calculated as the product of $E_R$ values for individual obstacles. Thus, the overall $E_R$ value for $O_s$-$O_1$ unlooped DNA equals 5.1. For $O_2$-$O_1$ unlooped DNA, $E_R$ equals 0.89 at the near ($O_2$) obstacle and 0.97 for the far ($O_1$) obstacle. Overall, the roadblock strength of unlooped $O_2$-$O_1$ DNA is approximately 1.

The λ CI repressor can also form looped and unlooped topologies. For λ CI-mediated looped DNA, $E_R$ equals 8.2. However, the actual roadblock strength is thought to be much less (see discussion in main text). For λ CI bound to unlooped DNA (2), $E_R$ equals 1.07 at the first obstacle (OL) and 1.42 at the second obstacle ($O_R$), $E_R$ = 1.42. The composite roadblock strength is 1.52.

The 186 CI repressor induced only unlooped topology with the minimum roadblock strength of 1.

| template | binding site | Pb- | Pa- | Pb+ | Pa+ | $E_R$ | composite $E_R$ |
|---|---|---|---|---|---|---|---|
| unlooped Os-O1 | | | | | | | |
| near | Os | 0.55 | 0.4 | 0.47 | 0.06 | 5.7 | 5.1 |
| far | O1 | 0.25 | 0.18 | 0.05 | 0.04 | 0.9 | |
| Looped Os-O1 | | | | | | | |
| near | Os | 0.55 | 0.4 | 0.2 | 0.02 | 7.3 | 7.3 |
| far | O1 | N/A | N/A | N/A | N/A | | |
| unlooped O2-O1 | | | | | | | |
| near | O2 | 0.55 | 0.4 | 0.55 | 0.45 | 0.9 | 0.9 |
| far | O1 | 0.25 | 0.18 | 0.27 | 0.2 | 1.0 | |
| Looped O2-O1 | | | | | | | |
| near | O2 | 0.55 | 0.4 | 0.4 | 0.02 | 14.5 | 14.5 |
| far | O1 | N/A | N/A | N/A | N/A | | |
| unlooped OL- OR | | | | | | | |
| near | OL | 0.61 | 0.5 | 0.38 | 0.29 | 1.1 | 1.5 |
| far | OR | 0.24 | 0.17 | 0.06 | 0.03 | 1.4 | |
| Looped OL- OR | | | | | | | |
| near | OL | 0.61 | 0.5 | 0.1 | 0.01 | 8.2* | 8.2* |
| far | OR | N/A | N/A | N/A | N/A | | |
| unlooped 186 CI | | | | | | | |
| | pR/pL | 0.64 | 0.42 | 0.66 | 0.43 | 1.0 | 1.0 |

Table S1. The probabilities of RNA progress straddling obstacle positions and the associated roadblock strengths calculated for measurements on templates bearing the the lac, lambda CI, and 186 CI repressors. * means this value is likely to be overestimated. See text for discussion.

### Supplementary figures

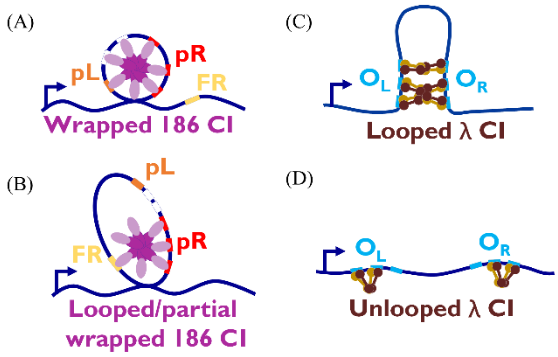


**Figure S1**. Different DNA binding structure of repressors. (A) The 186 CI wheel bound at pR/pL wraps DNA. (B) 186 CI binding at pR and FR forms a loop. (C) l CI bound at OL and OR mediate a DNA loop. (D) l CI dimers of dimers bound to the OL and OR regions.

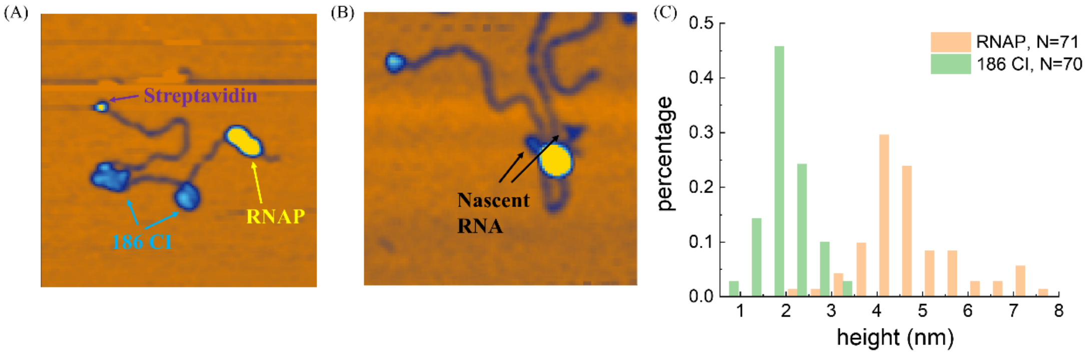


**Figure S2**. Height distinguishes RNAP from 186 CI. (A) Representative AFM image. (B) Nascent RNA can be found near RNAP (yellow) during transcription. Streptavidin label is cyan. (C) frequency distribution of the height of RNAP and 186 CI proteins measured by AFM

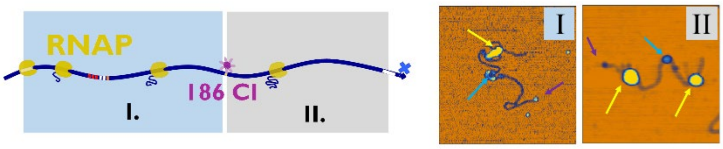


**Figure S3**. Transcription elongation complexes with 186 CI bound exclusively at the FR site. (Left) Cartoons describing the different regions where RNAP may be found with respect to the repressor bound to FR. (Right) AFM images of transcription elongation complexes along a DNA template bound by the 186 CI repressor at FR, color-coded as in the main text.

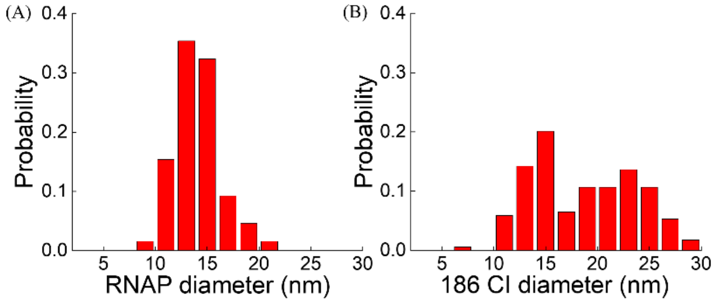


**Figure S4**. Comparison of RNAP (A) and 186 CI repressor (B) diameter distributions.

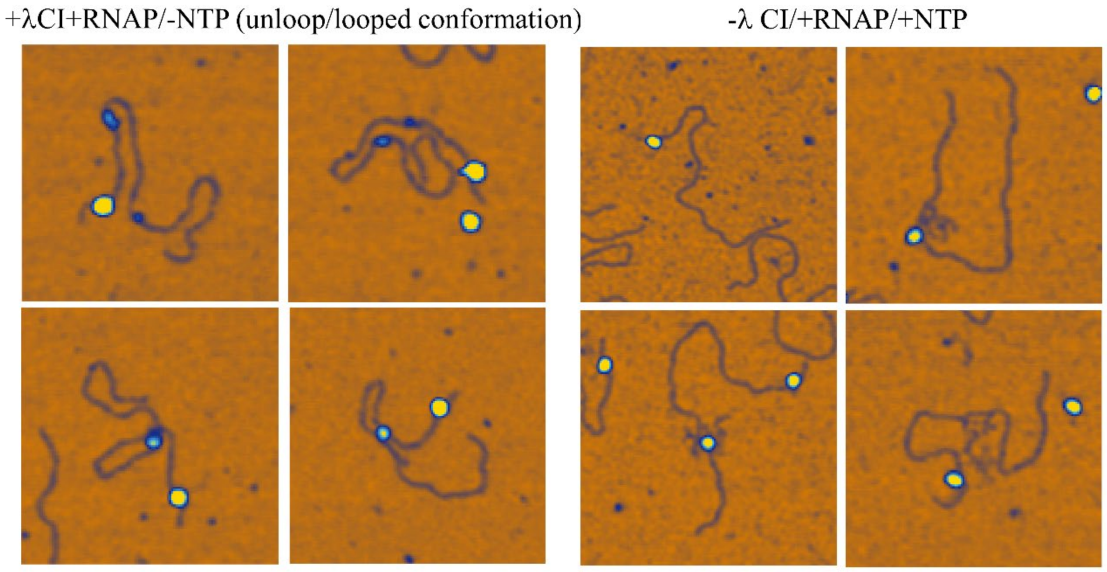


**Figure S5**. Gallery of AFM images of control experiments. (Left) RNAP is always found at the promoter in the absence of NTPs. (Right) In the absence of λ CI, RNAP is found at all stages of progress along the DNA template, as shown both by its position and the size of the nascent RNA associated with it.
